# Nearest-neighbor Projected-Distance Regression (NPDR) for detecting network interactions with adjustments for multiple tests and confounding

**DOI:** 10.1101/861492

**Authors:** Trang T. Le, Bryan A. Dawkins, Brett A. McKinney

## Abstract

Machine learning feature selection methods are needed to detect complex interaction-network effects in complicated modeling scenarios in high-dimensional data, such as GWAS, gene expression, eQTL, and structural/functional neuroimage studies for case-control or continuous outcomes. In addition, many machine learning methods have limited ability to address the issues of controlling false discoveries and adjusting for covariates. To address these challenges, we develop a new feature selection technique called Nearest-neighbor Projected-Distance Regression (NPDR) that calculates the importance of each predictor using generalized linear model (GLM) regression of distances between nearest-neighbor pairs projected onto the predictor dimension. NPDR captures the underlying interaction structure of data using nearest-neighbors in high dimensions, handles both dichotomous and continuous outcomes and predictor data types, statistically corrects for covariates, and permits statistical inference and penalized regression. We use realistic simulations with interactions and other effects to show that NPDR has better precision-recall than standard Relief-based feature selection and random forest importance, with the additional benefit of covariate adjustment and multiple testing correction. Using RNA-Seq data from a study of major depressive disorder (MDD), we show that NPDR with covariate adjustment removes spurious associations due to confounding. We apply NPDR to eQTL data to identify potentially interacting variants that regulate transcripts associated with MDD and demonstrate NPDR’s utility for GWAS and continuous outcomes.

## 1 Introduction

Pairwise epistasis is a measure of the effect of two genetic variants on a phenotype beyond what would be expected by their independent effects. There is evidence that these non-independent effects are pervasive (Breen *et al.*, 2012) and that higher-order interactions also play an important role in genetics (Weinreich *et al.*, 2013). A similar interaction effect can be observed in differential co-expression, where the phenotypic effect of one gene is modified d epending o n t he expression o f a nother g ene (Lareau *et al.*, 2015; De la Fuente, 2010). The embedding of these interactions in a regulatory network may lead to, not only pairwise interactions, but also higher-order epistasis network effects. Explicit modelling of these higher-order interactions would be computationally and statistically intractable (Riesselman *et al.*, 2018). Thus, computationally scalable feature selection methods are needed to capture these higher-order effects in high-dimensional data, such as genome-wide association (Arabnejad *et al.*, 2018) and RNA-Seq studies (Le *et al.*, 2018b).

Relief-based algorithms are efficient nearest-neighbor feature selection methods that are able to detect epistasis or statistical interaction effects in high-dimensional data without resorting to pairwise modeling of attributes (Urbanowicz *et al.*, 2018b; Kononenko *et al.*, 1997; McKinney *et al.*, 2009a; Robnik-Šikonja and Kononenko, 2003). We recently introduced the STatistical Inference Relief (STIR) formalism (Le *et al.*, 2018b) to address the lack of a statistical distribution for hypothesis testing and the related challenge of controlling the false positive rate of Relief-based scores. STIR extended Relief-based methods to compute statistical significance of attributes in dichotomous outcome data (e.g., case-control) by reformulating the Relief weight (McKinney *et al.*, 2013) as a pseudo t-test for the difference of means between projected-distances onto a given attribute. STIR is an effective approach with high power and low false-positive rate for data with main and interaction effects and is applicable to any predictor data type (continuous/gene expression or nominal/genetic variants). However, being based on a t-test, STIR does not apply to data with a continuous outcome (e.g., quantitative trait) and, like predecessor Relief-based methods, it does not correct for covariates.

In the current study, we introduce a new Nearest-neighbor Projected-Distance Regression (NPDR) approach that extends the STIR formalism to regression of nearest neighbors with the generalized linear model (GLM). The set of all nearest neighbor pairs is determined in the space of all attributes. Then for a given attribute, all pairs of neighbor instances are projected onto the attribute dimension and the one-dimensional distances (projected distances) become the observations in a GLM. The regression model of an attribute’s projected distances is linear for continuous outcomes and logistic for dichotomous outcomes, and either model may include corrections for covariates. The NPDR attribute importance score is given by its standardized regression coefficient and the statistical significance by its P value.

This NPDR framework also provides a natural way to control for covariates by including additional terms for the between-neighbor projected differences for each covariate within the GLM. Covariate adjustment is often neglected in machine learning, yet many biological and clinical omic studies involve potentially confounding covariates such as sex, bmi, and age (Le *et al.*, 2018a) or population stratification (Chen *et al.*, 2016). Some proposed methods for correcting machine learning algorithms include restricted permutation (Rao *et al.*, 2017), inverse probability weighting of training samples (Linn *et al.*, 2016) and penalized support vector machines (Li *et al.*, 2011). We demonstrate the effectiveness of NPDR to correct for confounding in an RNA-Seq study of major depressive disorder (MDD) in which there is a strong signal in the expression data due to the sex of study participants(Mostafavi *et al.*, 2014).

The flexible GLM formalism of NPDR opens nearest-neighbor feature selection to statistical inference for a broad class of problems. It can detect main effects and interactions for dichotomous and continuous outcome studies while adjusting for covariates and multiple hypothesis testing. The models allow any predictor data type such variants in GWAS or RNA-Seq expression levels. NPDR also improves attribute importance estimation compared to other Relief methods because NPDR includes the error or dispersion of the projected distances. The projected distance model gives the appearance of being univariate but implicitly accounts for interactions with all other attributes via the neighborhood calculation in the space of all attributes (omnigenic). Further, we extend the NPDR formalism to Lasso-like penalized feature selection.

The paper is organized as follows. In the Methods section, we develop the new formalism of NPDR to reformulate Relief-based scores as coefficients in a distanced-based GLM. We use the projected-distance regression formalism to implement a penalized version of NPDR. In the Results, we assess the performance of NPDR with simulations of main effects and network interactions, dichotomous outcomes with balanced and imbalanced classes, continuous outcomes, and continuous and genotypic predictors. We demonstrate improved precision and recall in comparison with standard Relief-based feature selection and random forest importance. We then apply NPDR to an RNA-Seq and GWAS study of MDD. From the RNA-Seq data, we show that NPDR removes spurious associations with MDD due to confounding by sex and identifies biologically relevant genes. Combining the RNA-Seq with the GWAS data, we perform an eQTL analysis with NPDR to identify potentially interacting variants that regulate transcripts associated with MDD and demonstrate NPDR’s utility for GWAS and continuous outcomes.

## 2 Materials and Methods

In this section, we develop the mathematical formalism needed to describe the projected distance regression in NPDR. We then construct the NPDR GLM models for common analysis situations, including continuous and dichotomous outcomes and adjustment for covariates. We also describe the simulation approach and real datasets for method validation.

### 2.1 Distance metrics and nearest neighbors

Because NPDR and other Relief-based feature selection methods are based on distances between instances, we first describe the algorithms and notation for identifying nearest neighbors in the space all attributes. We use the term attribute to refer to predictor variables, which may be continuous (e.g., expression) or categorical (e.g., variants). We use the term instance to refer to samples or subjects in a dataset.

#### 2.1.1 Distances and projections onto attributes

The distance between instances *i* and *j* in the data set *X*^*m*×*p*^ of *m* instances and *p* attributes is calculated in the space of all attributes (*a* ∈ *A*, |*A*| = *p*) using a metric such as

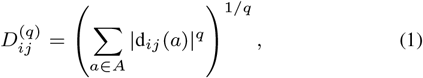

which is typically Manhattan (*q* = 1) in Relief but may also be Euclidean (*q* = 2). The quantity d_*ij*_(*a*), known as a “diff” in Relief literature, is the projection of the distance between instances *i* and *j* onto the attribute *a* dimension. The function d_*ij*_(*a*) supports any type of attribute (e.g., numeric/continuous versus categorical). For example, the projected difference between two instances *i* and *j* for a continuous numeric attribute, *a*, may be defined as

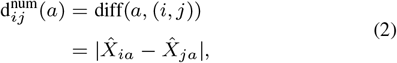

where *X̂* represents the standardized data matrix *X*. We use a simplified d_*ij*_(*a*) notation in place of the diff(*a*, (*i*, *j*)) notation that is customary in Relief-based methods. We also omit the division by max(*a*) − min(*a*) that is used by Relief to constrain scores to the interval from −1 to 1. As we show in subsequent sections, NPDR attribute importance scores are standardized regression coefficients with corresponding P values, so any scaling of the projected distances is unnecessary for comparing attribute scores. Thus, for the numeric-data projection, 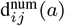, we simply use the absolute difference between row elements *i* and *j* of the data matrix *X*^*m*×*p*^ for the attribute column *a*.

The numeric projection function in Eq. (2) is appropriate for gene expression and other quantitative predictors and outcomes. For genome-wide association study (GWAS) data, where attributes are categorical, one simply modifies the type in the projection function from numeric to a discrete difference (Arabnejad *et al.*, 2018), but the projected-distance regression methods will be otherwise unchanged. The d_*ij*_(*a*) quantity is typically part of the metric to define the neighborhood, but it is also essential for computing the importance coefficients (Sec. 2.2.1). The projected-distance regression models below (Eqs. 8, 11, 14, and 15) will be fit for all nearest neighbors *i* and *j* in the defined neighborhood (discussed next in Eq. (3)).

#### 2.1.2 Nearest-neighbor ordered pairs

For NPDR, a single neighborhood is computed, regardless of whether the problem is classification or regression and regardless of whether a fixed-*k* or adaptive radius neighborhood method is used (Greene *et al.*, 2009; Urbanowicz *et al.*, 2018a; McKinney *et al.*, 2013). The NPDR neighbors are chosen blind to the outcome variable and then pairs of instances are assigned to hit or miss groups for dichotomous outcome data and assigned numeric differences for quantitative outcome data.

We define the NPDR neighborhood set 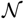 of ordered-pair indices as follows. Instance *i* is a point in *p* dimensions, and we designate the topological neighborhood of *i* as *N*_*i*_. This neighborhood is a set of other instances trained on the data *X*^*m*×*p*^ and depends on the type of Relief neighborhood method (e.g., fixed-*k* or adaptive radius) and the type of metric (e.g., Manhattan or Euclidean). If instance *j* is in the neighborhood of *i* (*j* ∈ *N*_*i*_), then the ordered pair is in the overall neighborhood (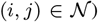 for the projected-distance regression analysis. The ordered pairs constituting the overall neighborhood can then be represented as nested sets:

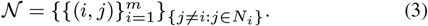

The cardinality of the set {*j* ≠ *i* : *j* ∈ *N*_*i*_} is *k*_*i*_, the number of nearest neighbors for subject *i*.

#### 2.1.3 Adaptive-radius and fixed-k Neighborhoods

The NPDR algorithm applies to any Relief neighborhood algorithm. In the NPDR analysis of real data, we use the multiSURF (Urbanowicz *et al.*, 2018a) adaptive-radius neighborhood, which uses a different radius for each instance. In the simulation anlysis, we use a fixed-*k* neighborhood that well-approximates mulitSURF for Relief and NPDR comparisons. The adaptive radius for an instance may be defined as the mean of its distances to all other instances subtracted by the fraction *α* of the standard deviation of this mean. More precisely, an instance *j* is in the adaptive *α*-radius neighborhood of 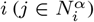 under the condition

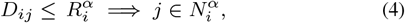

where the threshold radius for instance *i* is

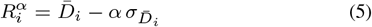

and

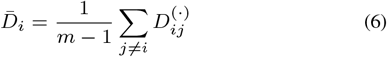

is the average of instance *i*’s pairwise distances (using Eq. 1) with standard deviation 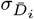. MultiSURF uses *α* = 1/2 (Granizo-Mackenzie and Moore, 2013).

Previously we showed empirically for balanced dichotomous outcome datasets that a good constant-*k* approximation to the expected number of neighbors within the multiSURF (*α* = 1/2) radii is *k* = *m*/6 (Le *et al.*, 2018b), where *m* is the number of samples. A more exact theoretical mean that shows the mathematical connection between fixed-*α* and fixed-*k* neighbor-finding methods is given by

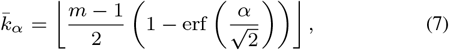

where we apply the floor to ensure the number of neighbors is integer. For data with balanced hits and misses in standard fixed-*k* Relief, one further divides this formula by 2, and then for multiSURF (*α* = 1/2), we find 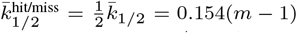, which is very close to our previous empirical estimate *m*/6. In the current study, when we compare multiSURF neighborhood methods with fixed-*k* neighborhoods, we use *k̅*_1/2_. Using this *α* = 1/2 value has been shown to give good feature selection performance by balancing power for main effects and interaction effects. However, the best value for *α* or *k* is likely data-specific and may be determined through nested cross-validation parameter tuning or other adaptive methods (McKinney *et al.*, 2013).

### 2.2 Nearest-neighbor Projected-Distance Regression (NPDR) with the generalized linear model

#### 2.2.1 Continuous outcomes: linear regression NPDR

Once the neighborhood 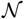 (Eq. 3) is determined by the distance matrix *D*_*ij*_ (Eq. 1) and the neighborhood method is chosen (e.g., fixed number of neighbors *k* or adaptive radius), we can compute the NPDR test statistic and P value for the association of an attribute with the phenotype. The NPDR model predictor vector is the attribute’s projected distances (d_*ij*_(*a*)) between all pairs of nearest-neighbor instances 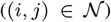. For continuous outcome data (quantitative phenotypes), the NPDR model outcome vector is the numeric difference (Eq. 2) between all nearest neighbors *i* and *j*. We find the parameters of the following model that minimize the least-squares error over 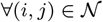:

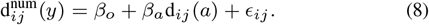

The 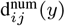 term on the left is the projected distance (diff) between instances *i* and *j* for a numeric phenotype *y* (Eq. 2) and *ϵ*_*ij*_ is the error term for this random variable. The predictor attribute *a* may be numeric or categorical, which determines the “type” used in the diff function on the right hand side of Eq.(8). The NPDR test statistic for attribute *a* is the *β*_*a*_ estimate with a one-sided hypothesis

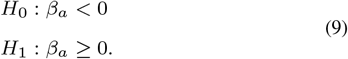

The *β*_*a*_ can be interpreted as the predicted change in the difference of the quantitative outcome between a pair of subjects when the projected distance of the attribute, *a*, changes by one unit. The attribute weights in the original RRelief algorithm (Robnik-Šikonja and Kononenko, 2003) can be described as a weighted covariance between the attribute neighbor projected distances, d_*ij*_(*a*), and the outcome neighbor differences, 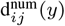. The extra weighting in RRelief is an exponentially decaying function of the rank of the distance between neighbors. The NPDR attribute weight is the standardized regression coefficient, 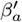, which is the covariance of the projected distances divided by the variance of the outcome projected distances. When the regression contains no additional covariates, the NPDR attribute weight can be written as the correlation between outcome and attribute neighbor projected distances:

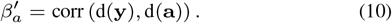

Thus, in the case of no covariates, NPDR for regression and RRelief have similar structure, but, as we show shortly, NPDR provides an improved attribute estimation, and the flexible NPDR framework can include additional sources of variation (i.e., adjust for confounding covariates).

#### 2.2.2 Linear regression NPDR with covariates

Previous Relief-based methods do not include the ability to adjust for covariates. The regression formalism of NPDR makes adding covariates straightforward. We simply compute the projected difference values 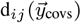 for the covariate attribute(s) between subjects on the neighborhood 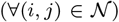 and include this as an additional projected distance term in the regression model:

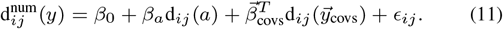

The above vector notation for the regression coefficients of the *p*_*c*_ covariates can be expanded as

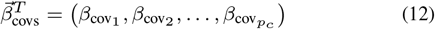

and the projection differences between instances *i* and *j* for each of the *p*_*c*_ covariates can be expanded as

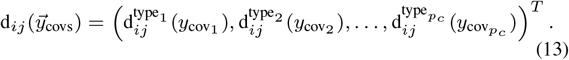

The superscripts in the projection operators above indicate the appropriate operator type for each covariate data type (e.g., numeric or categorical). In addition, the predictor attribute *a* may be numeric or categorical, which determines the type used in d_*ij*_(*a*). The NPDR test statistic is again *β*_*a*_ with alternative hypothesis *β*_*a*_ ≥ 0 as in Eq. (9), and the standardized 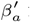 is the attribute importance score.

#### 2.2.3 Dichotomous outcomes with covariates: logistic regression NPDR

The STIR method was designed for statistical testing in dichotomous data, but as we have seen NPDR can also handle continuous outcomes and adjust for covariates. Here we show that the GLM formalism also enables NPDR to handle dichotomous outcome data (e.g., case-control phenotype). For dichotomous outcomes, NPDR models the probability 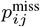 that subjects *i* and *j* are in the opposite class (misses) versus the same class (hits) from the neighbor projected distances with a logit function. We estimate the parameters of the following model for neighbors 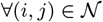:

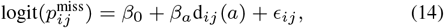

or if there are covariates,

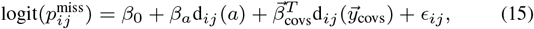

where 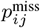 is the probability that subjects *i* and *j* have different phenotypes given the difference in their values for the attribute, *a*, and given the covariate differences. The outcome variable that is modeled by probability 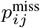 is a binary difference between subjects for the phenotype 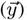:

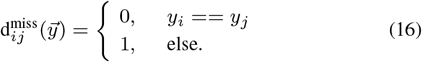

The *β*_*a*_ statistic can be interpreted in the following way. For a unit increase in the difference in the value of the attribute between two neighbors, we predict a change of 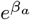 in the odds of the neighbors being in opposite classes. For dichotomous outcome data, we are interested in the alternative hypothesis that *β*_*a*_ > 0 because negative *β*_*a*_ values represent attributes that are irrelevant to classification. Thus, like NPDR linear regression, we are interested in testing one-sided hypotheses

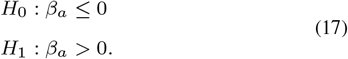

Nominal outcomes can be analyzed (similar to multi-state Relief-F) with NPDR by grouping all misses of an instance as one group.

#### 2.2.4 Regularized NPDR

One of the advantages of NPDR is its ability to estimate the confidence in the score of each attribute and provide a pseudo P value. This also allows NPDR to control for multiple testing by applying a false discovery rate adjustment to the collection of NPDR P values for all attributes. Here we develop a complementary feature selection method called regularized NPDR that combines all of the attribute difference vectors into one design matrix and constrains the coefficients to be non-negative, similar to the one-tailed test we use in standard NPDR. Specifically, we minimize the vector of regression coefficients, 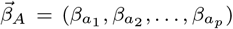, simultaneously for all attribute projections *a* ∈ *A*, subject to the coefficients being non-negative:

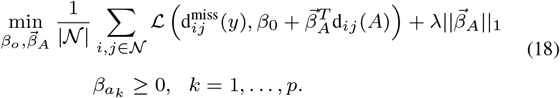

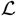 is the negative log-likelihood for each pair of instances *i* and *j* in neighborhood 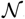, and d_*ij*_(*A*) represents the vector of diffs for fixed *i* and *j* for all attributes *a* ∈ *A*:

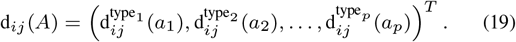

Our implementation uses a zero lower limit for the coefficients and the penalty strength λ > 0 is chosen by cross-validation (Zou and Hastie, 2005). For dichotomous outcomes, we use the binomial link function for the hit/miss projected distances in the likelihood optimization.

### 2.3 Properties of NPDR and existing Relief-based methods

Here we summarize the properties and capabilities of standard Relief-based methods (Urbanowicz *et al.*, 2018b) and the generalizations STIR (Le *et al.*, 2018b) and NPDR (Table 1). When there are no covariates for dichotomous outcome data, STIR (based on a pseudo t-test) and NPDR (based on regression with a logit model) are approximately equivalent given reasonable distribution assumptions (see Supplementary Fig. S1). However, by design, the NPDR framework is more flexible and able to handle covariate adjustment and continuous outcomes. In the current notation, the STIR null and alternative hypotheses would be

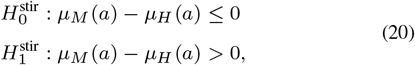

where

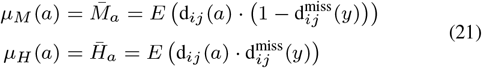

and 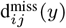 is given by Eq.(16). The STIR test statistic is a pseudo t-test (see Ref (Le *et al.*, 2018b)).

**Table 1.**
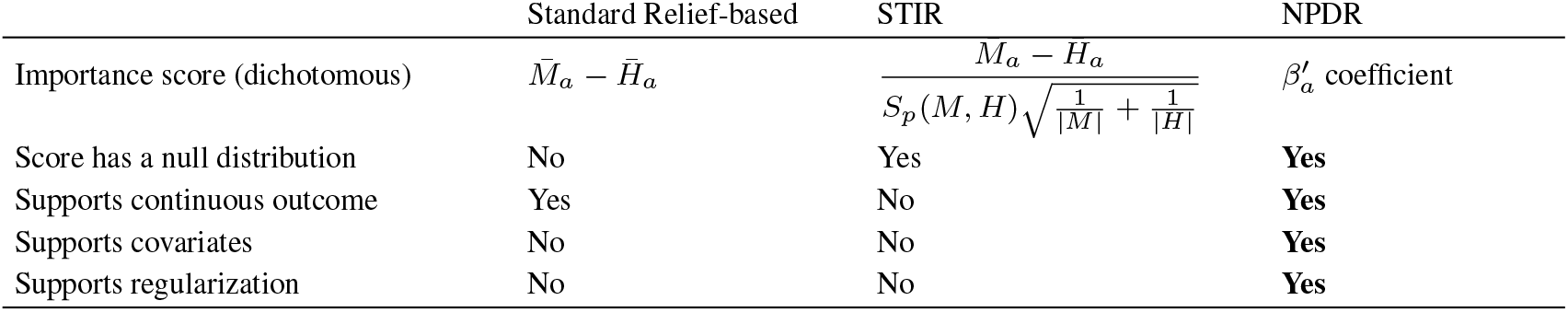
Properties of standard Relief-based methods and generalizations STIR and NPDR. The NPDR score is the standardized regression coefficient 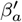 from a logistic model of projected distances (Eq. 14) for dichotomous outcomes and from a linear model (Eq. 8) for continuous outcomes. The quantity *S_p_* in STIR is the pooled standard deviation of the hit and miss means. Only the score for dichotomous (hit/miss) Relief is shown and STIR is limited to dichotomous outcomes (Le et al., 2018b).

For dichotomous outcomes, STIR improves attribute estimates over Relief weights (*M̅*_*a*_ – *H̅*_*a*_) by incorporating sample variance of the nearest neighbor distances in the denominator, which also enables STIR to estimate statistical significance with the assumptions of a t-test (Table 1). NPDR assumes intra- and inter-class differences are randomly sampled from one distribution and computes the importance score from a logistic regression 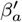. This regression-based generalization improves NPDR’s attribute estimates over Relief weights and enables statistical significance estimation for a wider range of problems than STIR.

### 2.4 Real and simulated datasets

#### 2.4.1 Simulation methods

To compare true positive and false positive performance for NPDR and other feature selection methods, we use the simulation tool from our private Evaporative Cooling (privateEC) software (Le *et al.*, 2017) that was designed to simulate realistic main effects, correlations, and interactions found in gene expression or resting-state fMRI correlation data. We simulate data with *m* = 200 subjects and *p* = 1, 000 real-valued attributes with 10% functional (true positive association with outcome). We simulate both balanced and imbalanced (75:25) dichotomous outcomes as well as continuous outcome data, and we simulate multiple main effects and interaction effects. We choose a sample size consistent with real gene expression data but on the smaller end to demonstrate a more challenging scenario. Likewise, the main effect size parameter (*b* = 0.8) was selected to be sufficiently challenging with power approximately 40% (Le *et al.*, 2017).

For interactions in dichotomous outcome data, we simulate network interactions using the differential co-expression network-based simulation tool in privateEC, which is described in Refs. (Le *et al.*, 2017; Lareau *et al.*, 2015). Briefly, first create a co-expression network on an Erdoős-Rényi random graph with 0.1 attachment probability, which is the critical value for a giant component. We give connected genes a higher average correlation, approximately *r*_connected_ = 0.8. This correlation between variables also controls the interaction effect size because we disrupt the correlation of target genes in cases but maintain correlation within controls, thereby creating a final d ifferential c orrelation n etwork. For GWAS simulations, we use benchmark simulated epistatic data from Ref. (Urbanowicz *et al.*, 2018a). Specifically, we use data with 20 variants (0.2 maximum minor allele frequency) and containing one pair of epistatic variants with a heritability of 10%. To make functional detection more challenging, we randomly select a small sample size of *m* = 200, including balanced and imbalanced (75 : 25) cases and control datasets.

We assess the feature-selection performance of each method by averaging the area under the recall curve (auRC) and the precision-recall curve (auPRC) across 100 replicate simulations for continuous predictors and 30 replicates for GWAS (genotypic) predictors. The auRC and auPRC metrics are good comparison tools for feature selection methods that, unlike NPDR, do not have a statistical significance threshold. The recall curve for a method is computed by sweeping across all score thresholds (from 0 to 100% selected) for a given dataset and calculating the recall (true positive rate) for each threshold. The auRC gives an overall measure of a method’s ability to detect functional attributes; however, in addition to detecting true positives, researchers are also interested in the false positive rate of a method. Thus, we also sweep across thresholds and compute the area under the precision versus recall curve (auPRC). The auPRC measures how well the method balances the ability to detect true positives versus including too many false positives across all thresholds

#### 2.4.2 RNA-Seq data and NPDR adjustment for confounding factors

To test the ability of NPDR to correct for confounding, we apply NPDR to the RNA-Seq study in Ref. (Mostafavi *et al.*, 2014) that consists of 15,231 genes for 463 MDD cases and 452 controls. Of the 915 subjects, 641 are female and 274 are male. The chi-square between MDD and sex is 25.75 (*p* = 3.89*e* − 7), and there are 485 genes significantly associated with sex. Thus, there is high risk for confounding effects due to sex differences. We apply NPDR with a multiSURF neighborhood and compute importance scores of all genes with and without sex as a covariate to isolate confounding genes. All resulting p-values (from STIR and NPDR) are adjusted for multiple testing. Attributes with adjusted p-values less than 0.05 are counted as a positive test (null hypothesis rejected), else the test is negative.

#### 2.4.3 eQTL data and NPDR for GWAS and quantitative outcomes

We perform an eQTL analysis with NPDR feature selection to identify potentially interacting variants that regulate transcripts associated with MDD and demonstrate NPDR’s utility for GWAS and continuous outcomes. The MDD RNA-Seq study described above includes 915 GWAS subjects genotyped with the Illumina Omni1-Quad microarray(Mostafavi *et al.*, 2014). We use NDPR to test for the cis- (1Mb from the gene’s transcription start site) and trans-eQTL influence on one of the gene expression levels associated with MDD (SCAI gene). We included 281,648 variants following GWAS filtering. We remove variants with a deviation from Hardy-Weinberg equilibrium (P < 0.0001 in controls) and a minor allele frequency (MAF) < 0.01, and we use linkage disequilibrium (LD) pruning to reduce the potential bias of correlation on interaction and distance calculations. SNPs are recursively removed within a sliding window along a given chromosome based on a pairwise LD of 0.5. We control for MDD status in the NPDR models to isolate more direct influence of variants on expression rather than MDD association. We use the Eq. (11) NPDR model and an allele mismatch operator for the SNP attribute projections (Arabnejad *et al.*, 2018).

### 2.5 Software availability

Detailed simulation and analysis code needed to reproduce the results in this study is available at https://github.com/lelaboratoire/npdr-paper (R version 3.5.0). The *npdr* R package is available at https://insilico.github.io/npdr/.

## 3 Results

### 3.1 Simulation results

Because Relief scores do not have a distribution for statistical significance, we compare score quality of methods by calculating the area under the recall (true positive rate) curve for a grid of score thresholds (Fig. 1). For simulated genotype data with epistasis (Fig. 1A), Relief and NPDR have similar recall for detecting the two interacting variants and higher recall than random forest. NPDR has a higher recall than Relief and random forest for simulated continuous-valued attributes with 100 network interactions in a background 1,000 total attributes (Fig. 1B). The performance of all methods decreases when there is imbalance (Fig. 1 right panels), but the relative performance between methods is the same.

**Fig. 1.**
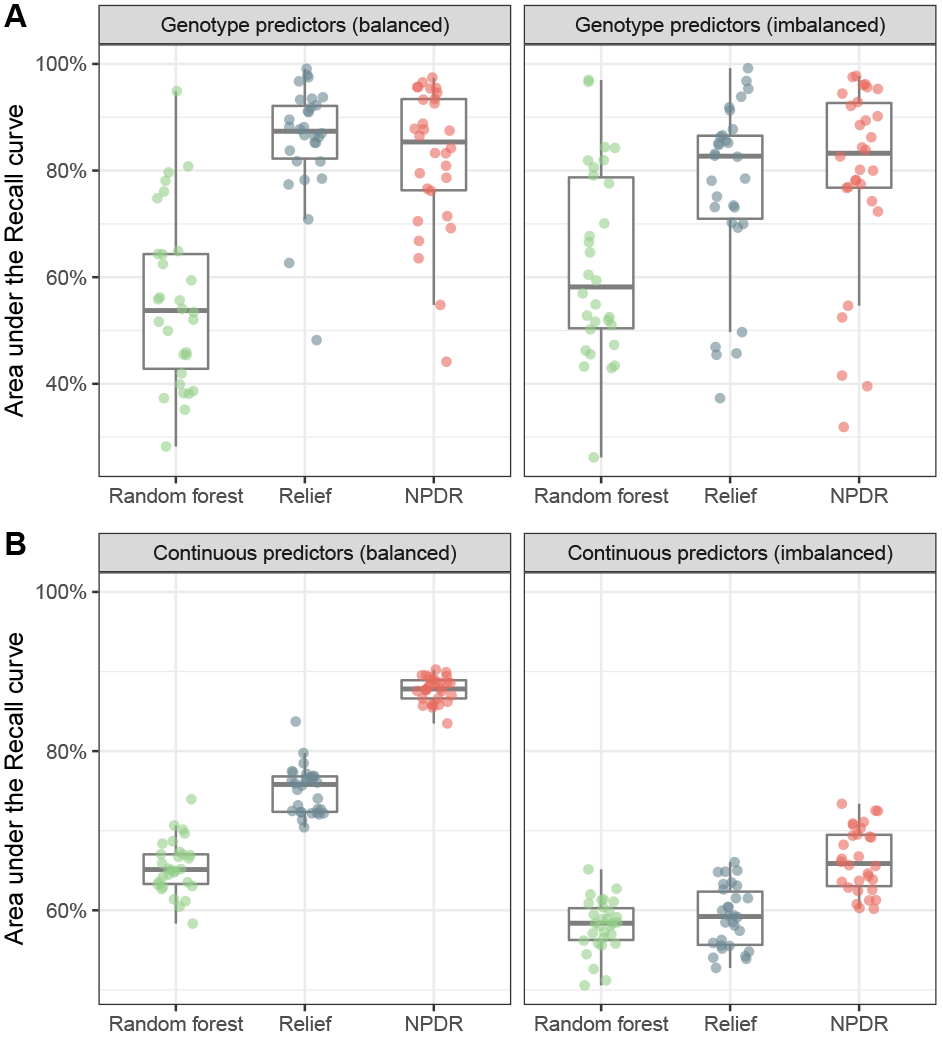
Recall (true positive rate) for detection of interaction effects in case-control data. For 30 replicate genotype simulations with epistasis (A, top row), the area under the recall curve (auRC) is compared for NPDR, Relief and random forest for balanced (left 50:50) and imbalanced (right 75:25) case-control data. The auRC measures the ability of each method to detect the pair of interacting variants with heritability 0.1 in *p* = 20 total attributes. For 100 replicate continuous-attribute simulations with a network of interactions (B, bottom row), the area under the recall curve (auRC) is compared for the three methods for balanced (left 50:50) and imbalanced (right 75:25) case-control data. The auRC measures the ability of each method to detect the 100 interacting attributes in *p* = 1, 000 total attributes. All simulations use *m* = 200 samples.

A high true positive rate is an important consideration, but this could come with a risk of a higher false positive rate, which can be measured by the feature selection precision. For one simulation with continuous-valued attributes, we illustrate the Precision-Recall Curve (PRC) for a grid of attribute importance thresholds (Fig. 2A) for each method. NPDR shows higher area under the PRC (auPRC) than Relief and random forest for continuous outcomes with main effects (left) and dichotomous outcomes with network interaction effects (right). Across 100 replicate simulations for each simulation type, NPDR shows significantly higher auPRC than random forest and Relief (both *P* < 0.0001, Fig. 2B).

**Fig. 2.**
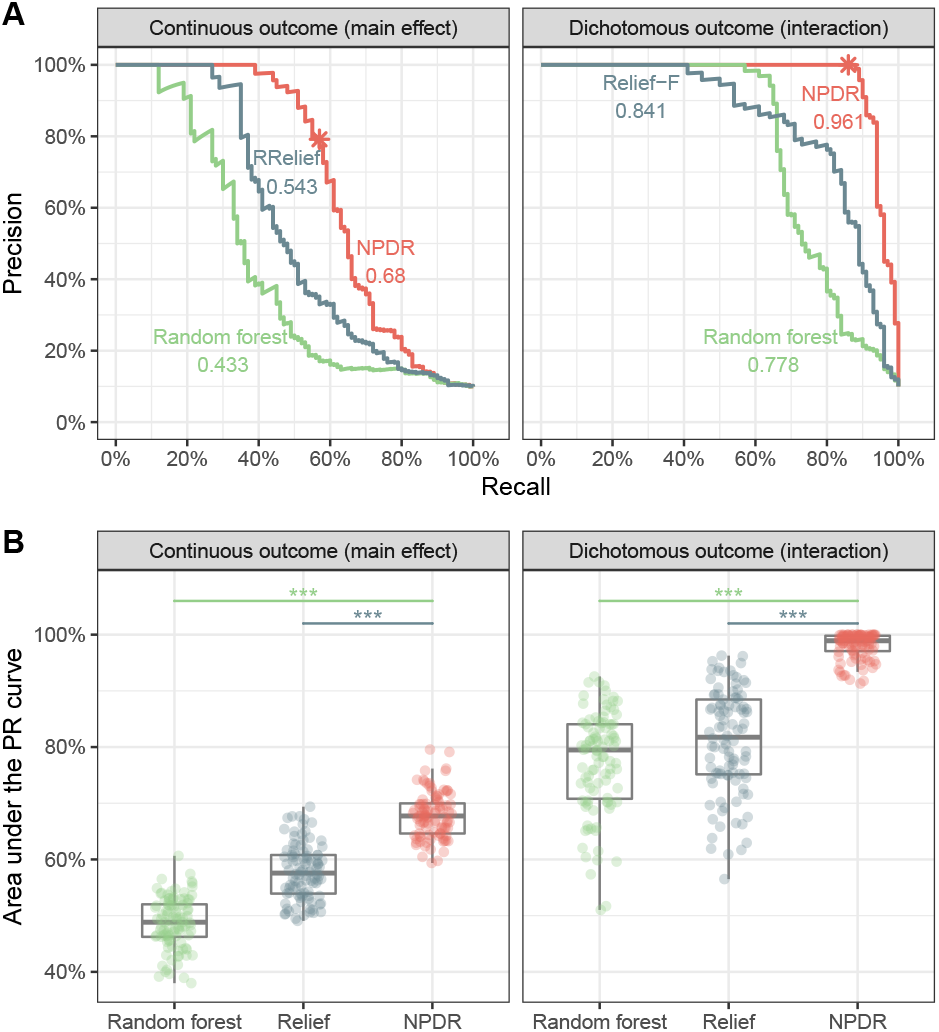
Precision-Recall for detection of simulated functional variables. For one replicate simulation (A, top row), precision-recall curves (PRC) are displayed for continuous outcome data with main effects (left) and dichotomous outcome data with interaction effects (right). The area under the PRC (auPRC) value is reported next to each method’s curve: NPDR, Relief-based and random forest. The ∗ indicates the NPDR 0.05 adjusted cutoffs (scores shown in Supplementary Fig. S2). For 100 replicate simulations (B, bottom row), the distributions of the auPRC values are compared for the methods. NPDR yields statistically significant higher auPRC than Relief or random forest (∗∗∗ indicate P < .0001). All simulations use *m* = 200 samples and *p* = 1, 000 attributes with 100 functional.

We use auPRC to compare other machine learning methods with NPDR because Relief and random forest lack a null distribution, whereas NPDR has an approximate distribution for hypothesis testing. NPDR correctly detects 57 out of 100 functional attributes in a continuous-outcome main effect simulation (Supplementary Fig. S2A) and 86 out of 100 functional attributes in a dichotomous-outcome interaction simulation (Supplementary Fig. S2B) using an adjusted P value threshold. Given any vertical score cutoff (Supplementary Fig. S2), it is difficult for Relief or random forest to detect most of the functional attributes without including many more false positives than NPDR. As shown by the auPRC (Fig. 2), NPDR includes fewer false positives than the other methods as it also detects more functional attributes.

NPDR importance scores are highly correlated with Relief-F scores with r = 0.869 for continuous outcome main effects and 0.848 for dichotomous outcome interaction effects (Supplementary Fig. S2). This correlation is expected because both methods are nearest-neighbor based. The correlation of NPDR with random forest scores is lower than with Relief-F, where r = 0.692 for continuous outcome main effects and 0.578 for dichotomous outcome interaction effects. The correlation for interaction effect simulations is lower than main effects because random forest underestimates the importance of interacting attributes as the attribute dimensionality becomes large compared to the number of functional attributes (McKinney et al., 2009b; Winham et al., 2012).

In addition to auRC and auPRC performance metrics, we also use the area under the Receiver Operating Characteristics curve (auROC), which also shows NPDR has statistically significant higher feature selection performance than random forest and Relief (both P < 0.0001, see Supplementary Fig. S3). We also compare NPDR (using a logistic model) with STIR (based on a t-test) for the dichotomous outcome data with interaction effects (Supplementary Fig. S1). Although standard logistic regression and the t-test have slightly different assumptions on the distribution of samples, NPDR and STIR yield highly correlated scores for dichotomous data with interaction effects, where the correlation between the P values produced from the two methods ranges from 0.9827 to 0.9994 in 100 replicate simulations.

In the simulations for continuous and dichotomous outcomes, respectively, we use NPDR with a linear model (Eq. 8) and logistic model (Eq. 14) to compare with standard RRelief and Relief-F (R package CORElearn) and with random forest regression and classification (permutation importance, R package randomforest). To provide a fair comparison of Relief and NPDR scores, we control for the neighborhood method by using the same fixed-*k* neighborhood 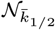 for all methods The value *k̅*_1/2_ = 30 (Eq. 7) is the expected number of nearest neighbors corresponding to a multiSURF neighborhood for the simulated sample size *m* = 200. In the simulation analysis, this k value is used for NPDR nearest neighbors, and one half this value is used for ReliefF because its k represents nearest hits and misses separately. For a dataset of the size simulated in this study (*m* = 200 samples and *p* = 1, 000 attributes with 100 functional), on a desktop with an Intel Xeon W-2104 CPU and 32GB of RAM, NPDR has a 24-second and 3-second runtime for dichotomous and continuous outcome data, respectively.

### 3.2 RNA-Seq NPDR analysis for MDD with confounding

NPDR with covariate adjustment effectively removes sex-related confounding genes in the RNA-Seq study of MDD in Ref. (Mostafavi *et al.*, 2014). We apply NPDR with the multiSURF neighborhood 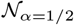 and an adjustment for the sex covariate (Eq. 15). This study contains numerous genes that are potentially confounded by sex differences. The sex variable is significantly associated with MDD, and 485 out of the 15,231 genes are associated with sex (Bonferroni-adjusted P value < 0.0001). The NPDR adjustment removes the genes that are most likely spurious MDD associations due to confounding (dark points below the horizontal 0.05 adjusted significance line in Fig. 3) compared to NPDR without adjustment. Not only do these removed genes have strong differential expression based on sex, but many of these genes, such as PRKY, UTY, and USP9Y, are Y-linked and mainly expressed in testis. For example, the RPS4Y2 ribosomal protein S4 Y-linked 2 has been shown by tissue specific studies to mainly express in prostate and testis (Lopes et al., 2010) while RPS4X (also associated with sex in the data) is most expressed in the ovary. The NPDR runtime for this RNA-Seq dataset (*m* = 915 samples and *p* = 15, 231 attributes) was approximately 2.3 hours on a desktop with an Intel Xeon W-2104 CPU and 32GB of RAM.

**Fig. 3.**
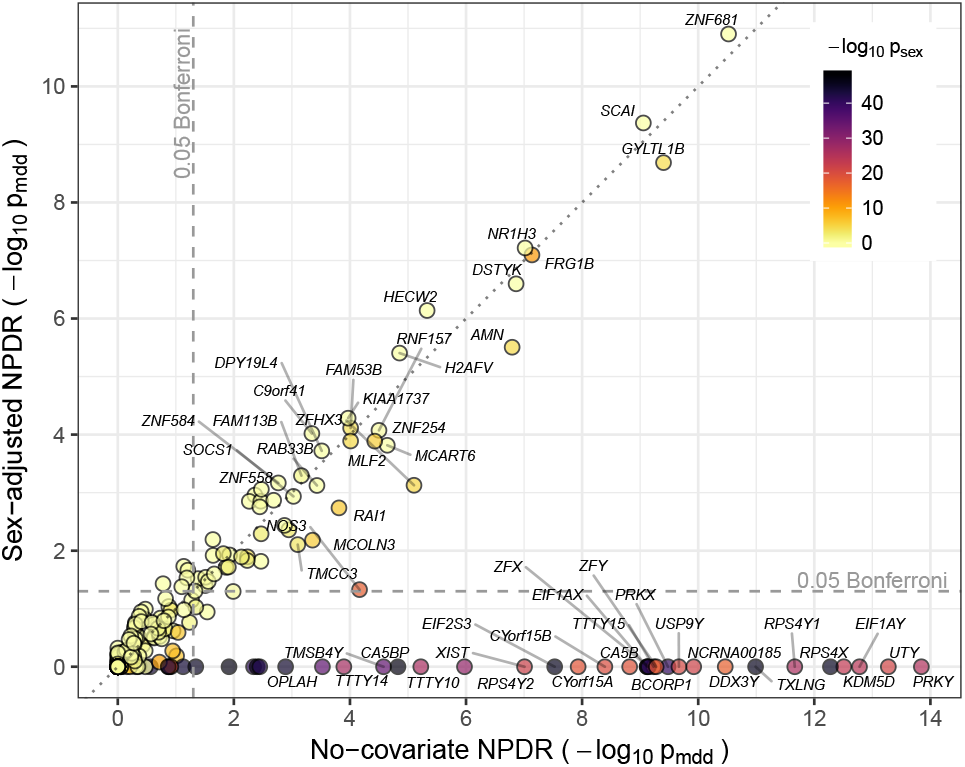
Comparison of NPDR major depressive disorder associations with and without covariate adjustment. Gene scatter plot of −log_10_ significance using NPDR without correction for sex (horizontal axis) and with correction for sex (vertical axis). Genes with adjusted *p*_mdd_ < 0.001 by either method are labeled. NPDR without sex correction finds 87 genes associated with MDD at the Bonferroni-adjusted 0.05 level (right of vertical dashed line), 53 of which are also significantly correlated with sex (adjusted *p*_sex_ < 0.05). NPDR with adjustment for sex finds 56 genes associated with MDD at the Bonferroni-adjusted 0.05 level (above horizontal dashed line), 19 of which are significantly correlated with sex. The most highly associated genes with sex are eliminated by adjustment (dark genes below the horizontal dashed line) but remain in the non-adjusted set (right of dashed vertical line).

### 3.3 eQTL analysis using NPDR with GWAS and quantitative outcomes

We perform an eQTL analysis with NPDR to demonstrate its ability to analyze continuous outcomes and SNP predictors from GWAS. We choose to test for eQTLs that influence the expression of SCAI (suppressor of cancer cell invasion, involved in cell migration and regulation of cell cycle on chromosome 9), which was one of the stronger NPDR associations with MDD (Fig. 3) and was found previously to have a modest cis-eQTL effect in this dataset (Mostafavi *et al.*, 2014). One of the eQTLs found by NPDR (rs10997355) is an intron variant in CTNNA3 (catenin alpha-3 mediates cell–cell adhesion) on chromosome 10. In a study of schizophrenia, an intron variant near rs10997355 showed an interaction with maternal cytomegalovirus (CMV) status (Børglum *et al.*, 2014). Other studies of CTNNA3 and its nested gene LRRTM3 (encoding the Leucine-rich repeat transmembrane neuronal protein 3) have found associations with Alzheimer’s disease (Miyashita *et al.*, 2007) and autism spectrum disorder (Wang *et al.*, 2009). The top 100 NDPR eQTLs for SCAI are provided as supplementary information. The NPDR runtime was 7.06 hours (on a high performance computing environment with an Intel Xeon E5-2650v3 CPU, 20 nodes and 32GB of RAM).

## 4 Discussion

NPDR is the first method to our knowledge to combine projected distances and nearest-neighbors into a generalized linear model (GLM) regression framework to perform feature selection. The use of nearest neighbors enables the detection of interacting attributes, which NPDR shares with Relief-based methods. NPDR extends Relief in the following ways. (1) For feature selection with dichotomous outcomes, NPDR uses a logistic model to fit pairwise projected distance predictors onto hit and miss class differences. (2) This regression formalism provides a simple mechanism for the projected distance-based NPDR to correct for covariates, which is often neglected in machine learning and has been a limitation of Relief-based methods. (3) For continuous outcome data, NPDR simply performs a linear regression between the outcome and attribute projected distances between neighbors. The NPDR regression coefficient is an effective importance score and has an interpretation of variation explained in the outcome variable distances between subjects. (4) For any outcome data type (dichotomous or continuous) and any predictor data type, the NPDR score of an attribute includes the error of the estimate via the standardized regression coefficient, which enables statistical inference, significance testing of attribute scores, and adjustment for multiple testing. This additional variation is neglected by Relief-based methods. (5) We introduced a regularized NPDR that adds another layer of multivariate modeling to an already multi-dimensional nearest-neighbor method to shrink correlated projected attribute differences.

We assessed NPDR’s true positive rate and ability to control false positives using realistic simulations with main effects, network interactions, continuous outcomes, dichotomous outcomes with balanced and imbalanced classes, and categorical (genotypes) and continuous (expression) predictors. We showed that the statistical performance using NPDR P values is the same as STIR, where STIR is limited to dichotomous outcome data. In other words, by modeling hit/miss differences between neighbors with a logit link, NPDR can be used safely instead of STIR with the added benefit of covariate correction and the analysis of quantitative traits. The simulations provided empirical evidence for the improved recall and precision of NPDR over Relief-based and Random Forest feature selection.

A related distance-based regression method is Multivariate Distance Matrix Regression (MDMR) (Schork and Zapala, 2012). The MDMR approach uses an F-statistic to test for the association of distance matrices between two sets of factors. The MDMR regression is performed for the distance matrix for all pairs of instances, not a subset of nearest neighbors like NPDR, which makes MDMR susceptible to missing interactions. The use of local neighborhoods allows NPDR to remove imposter/irrelevant instances from the neighborhood and detect interactions in the higher dimensional space. Another distinction between methods is that NPDR projects distances onto each attribute, allowing for hypothesis testing of individual attributes (i.e., perform feature selection), whereas MDMR uses specified sets of attributes. NPDR uses the context of all attributes to compute nearest neighbors, but it focuses on the projected regression of each attribute at a time and uses the nearest neighbors to allow for detection of interactions. However, NPDR can also be used to compute the importance of sets of factors, similar to MDMR. An example of this is the penalized version of NPDR that uses the set of all attributes in a nearest-neighbor projected-distance multiple regression.

The NPDR implementation can use fixed-*k*, fixed radius, or adaptive radius neighborhood methods. To provide a fair comparison between Relief and NDPR methods, we used the same fixed-*k* for all simulation analyses. Power for detecting main effects is highest with the myopic maximum 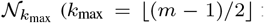 for balanced data). However, real biological data will likely contain a mixture of main effects and interaction network effects (McKinney and Pajewski, 2012). Thus, for all simulation analyses we used a fixed-k that has been shown to balance main effects and interactions (Eq. 7 with *α* = 1/2) (Le *et al.*, 2018b). NPDR feature selection can be embedded in the backwards elimination of private Evaporative Cooling (privateEC) or in a nested cross-validation for feature selection and classification (Le *et al.*, 2017) or tuning *α* or *k* to balance main effect and interaction detection.

The ability to incorporate covariates into NPDR models addresses the important but often neglected issue of confounding factors in machine learning. We applied NPDR to a real RNA-Seq dataset for MDD to demonstrate the identification of biologically relevant genes and the removal of spurious associations by covariate correction. NPDR with sex as a covariate adjustment successfully removed X and Y-linked genes and genes highly expressed in sex organs. However, it is important to note that some genes removed due to a shared association with sex may be important for the pathophysiology of MDD or for classification accuracy. Thus, covariate adjustment in NPDR is a useful option to inform a holistic analysis of a given dataset.

Application to GWAS data required no additional modifications of the algorithm other than specification of a different diff projection operator for categorical variables (Arabnejad *et al.*, 2018), and the covariate option allows principal components to be included to adjust for population structure. One of the trans-eQTLs found by NPDR in CTNNA3 (rs10997355) for SCAI may suggest a gene-environment interaction due to maternal CMV infection. In addition, it has been suggested that exposure to CMV may increase mood disorder risk through interactions with susceptibility variants (Kim *et al.*, 2007).

To expand variant discovery and demonstrate the ability of NDPR to analyze large GWAS data, we performed a conservative LD pruning that removes some correlation between variants while still leaving a substantial number of informative variants. Our simulations for continuous predictors included correlation as well as interactions, but because of the omni-genic nature of NPDR, further investigation is needed to understand the effect of correlation on the relative ranking of variants and the effect on nearest-neighbor calculations. A challenge in NPDR analysis is the inherent dependence between neighbors in the models, which violates distribution assumptions of regression and leads to artificially lower P-values. Our results indicate that the Bonferroni procedure effectively controls type I error despite deflated P-values. However, future studies are needed to investigate strategies to account for dependence between neighbor-pair observations

## Supporting information

Complete Supplement

## References

Arabnejad, M., Dawkins, B., Bush, W., White, B., Harkness, A., and McKinney, B. A. (2018). Transition-transversion encoding and genetic relationship metric in relieff feature selection improves pathway enrichment in gwas. BioData mining,11, 23.

Børglum, A., Demontis, D., Grove, J., Pallesen, J., Hollegaard, M. V., Pedersen, C., Hedemand, A., Mattheisen, M., Uitterlinden, A., Nyegaard, M., et al. (2014). Genome-wide study of association and interaction with maternal cytomegalovirus infection suggests new schizophrenia loci. Molecular psychiatry, 19(3), 325.

Breen, M., Kemena, C., Vlasov, P., Notredame, C., and Fa, K. (2012). Epistasis as the primary factor in molecular evolution. Nature, 490(7421), 535–8.

Chen, H., Wang, C., Conomos, M. P., Stilp, A. M., Li, Z., Sofer, T., Szpiro, A. A., Chen, W., Brehm, J. M., Celedón, J. C., et al. (2016). Control for population structure and relatedness for binary traits in genetic association studies via logistic mixed models. The American Journal of Human Genetics, 98(4), 653–666.

De la Fuente, A. (2010). From differential expression to differential networking– identification of dysfunctional regulatory networks in diseases. Trends in genetics, 26(7), 326–333.

Granizo-Mackenzie, D. and Moore, J. H. (2013). Multiple threshold spatially uniform relieff for the genetic analysis of complex human diseases. In European Conference on Evolutionary Computation, Machine Learning and Data Mining in Bioinformatics, pages 1–10. Springer.

Greene, C. S., Penrod, N. M., Kiralis, J., and Moore, J. H. (2009). Spatially Uniform ReliefF (SURF) for computationally-efficient filtering of gene-gene interactions. BioData Mining, 2, 5.

Kim, J. J., Shirts, B. H., Dayal, M., Bacanu, S.-a., Wood, J., Xie, W., Zhang, X., Chowdari, K. V., Yolken, R., Devlin, B., et al. (2007). Are exposure to cytomegalovirus and genetic variation on chromosome 6p joint risk factors for schizophrenia? Annals of medicine, 39(2), 145–153.

Kononenko, I., Šimec, E., and Robnik-Šikonja, M. (1997). Overcoming the Myopia of Inductive Learning Algorithms with RELIEFF. Applied Intelligence, 7(1), 39–55.

Lareau, C. A., White, B. C., Oberg, A. L., and McKinney, B. A. (2015). Differential co-expression network centrality and machine learning feature selection for identifying susceptibility hubs in networks with scale-free structure. BioData mining, 8(1), 5.

Le, T. T., Simmons, W. K., Misaki, M., Bodurka, J., White, B. C., Savitz, J., and McKinney, B. A. (2017). Differential privacy-based evaporative cooling feature selection and classification with relief-f and random forests. Bioinformatics, 33(18), 2906–2913.

Le, T. T., Kuplicki, R. T., McKinney, B. A., Yeh, H.-w., Thompson, W. K., and Paulus, M. P. (2018a). A nonlinear simulation framework supports adjusting for age when analyzing brainage. Frontiers in aging neuroscience, 10.

Le, T. T., Urbanowicz, R. J., Moore, J. H., and McKinney, B. A. (2018b). Statistical inference relief (stir) feature selection. Bioinformatics.

Li, L., Rakitsch, B., and Borgwardt, K. (2011). ccsvm: correcting support vector machines for confounding factors in biological data classification. Bioinformatics, 27(13), i342–i348.

Linn, K. A., Gaonkar, B., Doshi, J., Davatzikos, C., and Shinohara, R. T. (2016). Addressing confounding in predictive models with an application to neuroimaging. The international journal of biostatistics, 12(1), 31–44.

Lopes, A. M., Miguel, R. N., Sargent, C. A., Ellis, P. J., Amorim, A., and Affara, N. A. (2010). The human rps4 paralogue on yq11. 223 encodes a structurally conserved ribosomal protein and is preferentially expressed during spermatogenesis. BMC molecular biology, 11(1), 33.

McKinney, B. and Pajewski, N. (2012). Six degrees of epistasis: statistical network models for gwas. Frontiers in genetics, 2, 109.

McKinney, B. A., Crowe, J. E., Guo, J., and Tian, D. (2009a). Capturing the spectrum of interaction effects in genetic association studies by simulated evaporative cooling network analysis. PLoS genetics, 5(3), e1000432.

McKinney, B. A., Crowe Jr, J. E., Guo, J., and Tian, D. (2009b). Capturing the spectrum of interaction effects in genetic association studies by simulated evaporative cooling network analysis. PLoS genetics, 5(3), e1000432.

McKinney, B. A., White, B. C., Grill, D. E., Li, P. W., Kennedy, R. B., Poland, G. A., and Oberg, A. L. (2013). Reliefseq: a gene-wise adaptive-k nearest-neighbor feature selection tool for finding gene-gene interactions and main effects in mrna-seq gene expression data. PLoS one, 8(12), e81527.

Miyashita, A., Arai, H., Asada, T., Imagawa, M., Matsubara, E., Shoji, M., Higuchi, S., Urakami, K., Kakita, A., Takahashi, H., et al. (2007). Genetic association of ctnna3 with late-onset alzheimer’s disease in females. Human molecular genetics, 16(23), 2854–2869.

Mostafavi, S., Battle, A., Zhu, X., Potash, J. B., Weissman, M. M., Shi, J., Beckman, K., Haudenschild, C., McCormick, C., Mei, R., et al. (2014). Type i interferon signaling genes in recurrent major depression: increased expression detected by whole-blood rna sequencing. Molecular psychiatry, 19(12), 1267.

Rao, A., Monteiro, J. M., Mourao-Miranda, J., Initiative, A. D., et al. (2017). Predictive modelling using neuroimaging data in the presence of confounds. NeuroImage, 150, 23–49.

Riesselman, A. J., Ingraham, J. B., and Marks, D. S. (2018). Deep generative models of genetic variation capture the effects of mutations. Nature methods, 15(10), 816–822.

Robnik-Šikonja, M. and Kononenko, I. (2003). Theoretical and empirical analysis of relieff and rrelieff. Machine learning, 53(1-2), 23–69.

Schork, N. J. and Zapala, M. A. (2012). Statistical properties of multivariate distance matrix regression for high-dimensional data analysis. Frontiers in genetics, 3, 190.

Urbanowicz, R. J., Olson, R. S., Schmitt, P., Meeker, M., and Moore, J. H. (2018a). Benchmarking relief-based feature selection methods for bioinformatics data mining. Journal of Biomedical Informatics, 85, 168–188.

Urbanowicz, R. J., Meeker, M., Cava, W. L., Olson, R. S., and Moore, J. H. (2018b). Relief-based feature selection: Introduction and review. Journal of Biomedical Informatics.

Wang, K., Zhang, H., Ma, D., Bucan, M., Glessner, J. T., Abrahams, B. S., Salyakina, D., Imielinski, M., Bradfield, J. P., Sleiman, P. M., et al. (2009). Common genetic variants on 5p14. 1 associate with autism spectrum disorders. Nature, 459(7246), 528.

Weinreich, D., Lan, Y., Wylie, C., and Heckendorn, R. (2013). Should evolutionary geneticists worry about higher-order epistasis? Curr Opin Genet Dev., 23(6), 700–7.

Winham, S. J., Colby, C. L., Freimuth, R. R., Wang, X., De Andrade, M., Huebner, M., and Biernacka, J. M. (2012). Snp interaction detection with random forests in high-dimensional genetic data. BMC bioinformatics, 13(1), 164.

Zou, H. and Hastie, T. (2005). Regularization and variable selection via the elastic net. Journal of the Royal Statistical Society: Series B (Statistical Methodology), 67(2), 301–320.

