## Supplementary material for "Nearest-neighbor Projected-Distance Regression (NPDR) for detecting network interactions with adjustments for multiple tests and confounding": Complete Supplement

### NPDR Supplementary Material

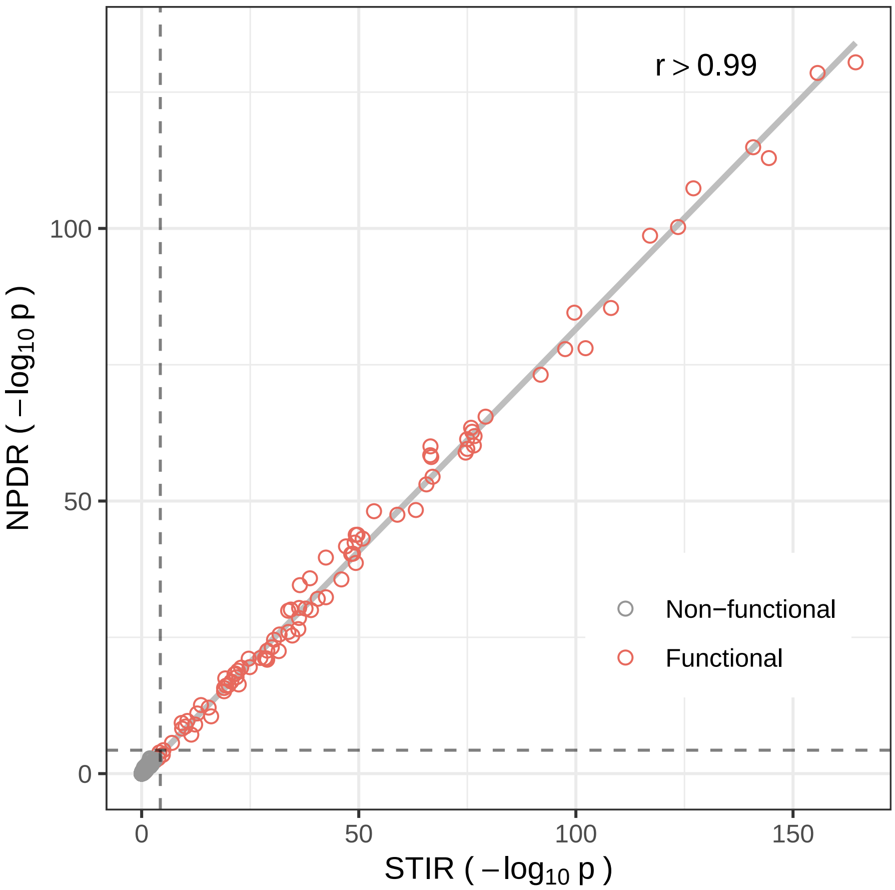


Figure S1: **Similarity between NPDR and STIR (dichotomous outcomes).** Comparison of -$\log_{10}$ P values for one interaction simulation of $m = 200$ (100 cases and 100 controls) and $p = 1000$ attributes with 100 functional. In 100 replicate simulations, correlation, $r$, between the two methods ranges from 0.9827 to 0.9994. STIR is based on a t-test of projected distances and NPDR is based on a logistic regression of projected distances. NPDR has the added benefit of handling continuous outcomes and covariate correction.

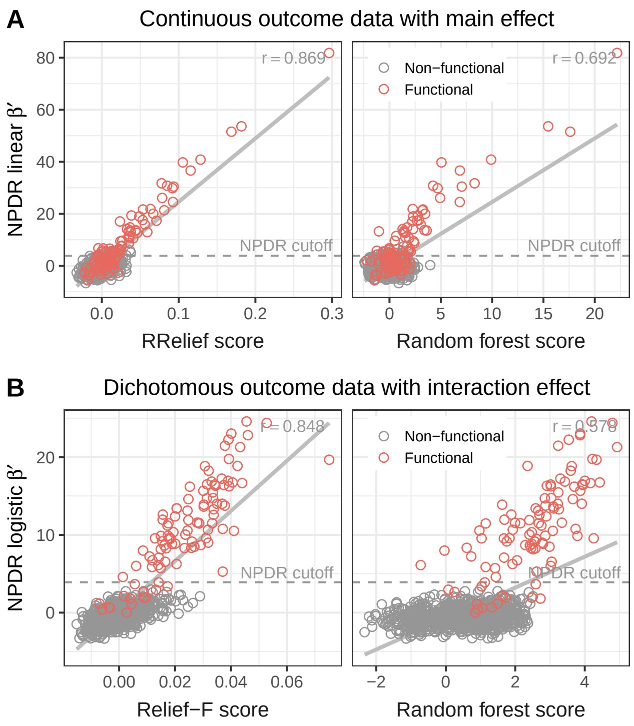


Figure S2: **Simulation comparison of importance scores.** Scatter plots of NPDR versus Relief-based scores (left) and NPDR versus random forest scores (right) for representative simulations of continuous outcome data with main effects (A, top row) and dichotomous outcome data with interaction effects (B, bottom row). Simulations use $m = 200$ samples and $p = 1,000$ attributes with 100 functional (orange). For continuous outcome (A), importance scores computed by RRelief weight, random forest percent increase in MSE and NPDR standardized linear regression coefficient ($\beta'$ from Eq. 10, main text). For dichotomous outcome (B), scores computed by Relief-F, random forest mean decrease in accuracy and NPDR standardized logistic regression coefficient ($\beta'$ from Eq. 14, main text). A regression line between the scores with correlation $r$ is displayed, and a 0.05 Bonferroni-adjusted cutoff (dashed) is shown for NPDR scores. There is no statistical threshold for Relief-based methods or random forest (area under the precision-recall curve (auPRC) is used to compare algorithm performance, see Fig. 1, main text).

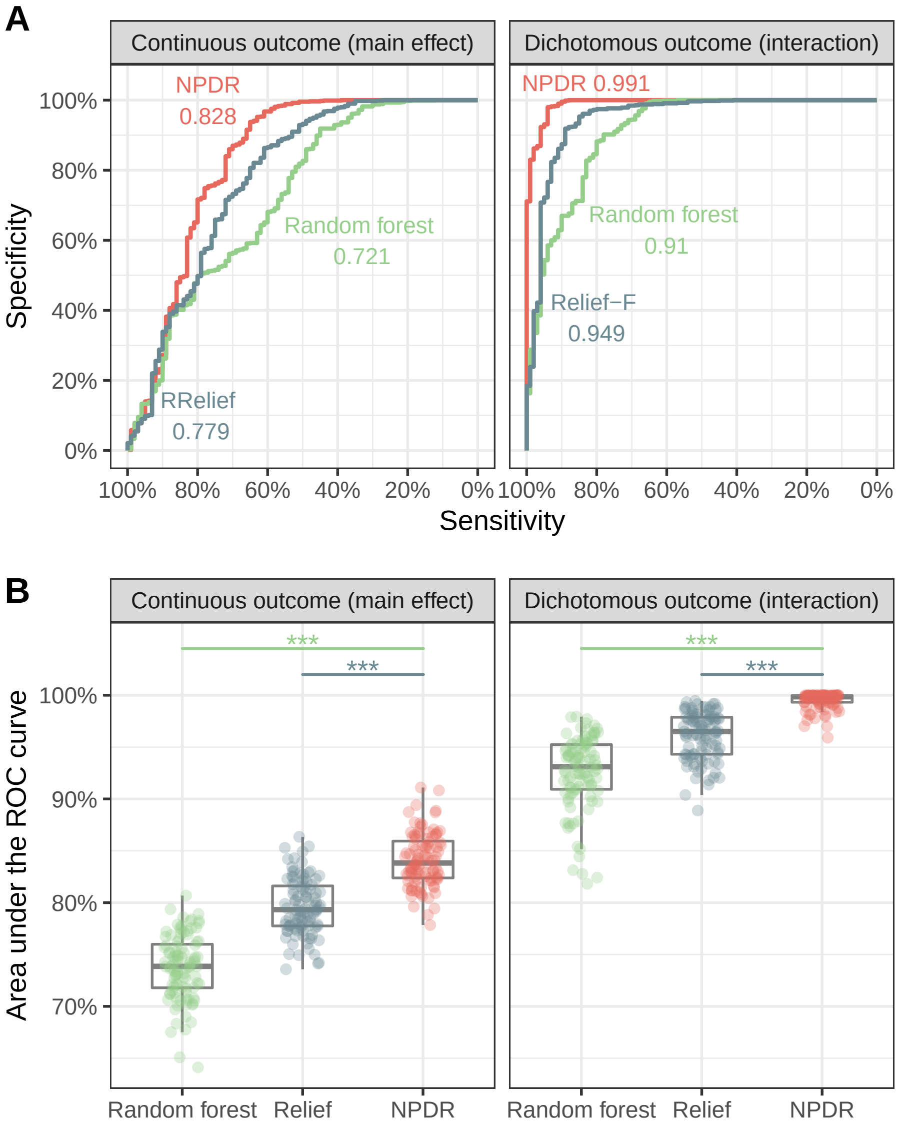


Figure S3: **Receiver Operating Characteristic (ROC) curves for Relief, NPDR and random forest feature selection.** For one replicate simulation (A), ROCs are displayed for continuous outcome data with main effects (left) and dichotomous outcome data with interaction effects (right). The auROC value is given for each method. For 100 replicate simulations of both simulation types (B), NPDR yields statistically significant higher auROC than Relief or random forest ($***$ indicate P $<$ .0001). All simulations use $m = 200$ samples and $p = 1,000$ attributes with 100 functional.

| NPDR Rank | rs-num | Chromosome | Ensembl Gene IDs | Missense Variant | Synonymous Variant | 5 Prime UTR Variant | 3 Prime UTR Variant | Non-coding Transcript Exon Variant | Intron Variant | NMD Transcript Variant | Non-coding Transcript Variant | Upstream Gene Variant | Downstream Gene Variant |
|---|---|---|---|---|---|---|---|---|---|---|---|---|---|
| 1 | rs4588246 | 2 | RN7SKP141 | | | | | | | | | | X |
| 2 | rs10937067 | 3 | AC007547.1 | | | | | | | | | X | |
| 3 | rs13170066 | 5 | MAST4 | | | | | | X | | X | | |
| 4 | rs4613744 | 5 | MIR583HG,AC104123.1 | | | | | | X | | X | | |
| 5 | rs16901512 | 8 | FAM84B,PCAT1 | | | | | | X | | X | X | |
| 6 | rs10997355 | 10 | CTNNA3 | | | | | | X | | | | |
| 7 | rs11061894 | 12 | WNT5B | | | | | | X | | | X | X |
| 8 | rs16909912 | 12 | PLBD1 | | | | | | X | | | | X |
| 9 | rs1259744 | 12 | AC023050.3,LINC02422 | | | | | | | | | X | |
| 10 | rs12322049 | 12 | | | | | | | | | | | |
| 11 | rs7297446 | 12 | TESC | | | | | | X | X | X | | |
| 12 | rs9319336 | 13 | | | | | | | | | | | |
| 13 | rs17711848 | 15 | THSD4,THSD4-AS1 | | | | | | X | | X | | |
| 14 | rs4312323 | 16 | COX6A2,ITGAD,AC026471.5 | | | | | | X | | X | X | X |
| 15 | rs16964218 | 16 | HMGB3P32 | | | | | | | | | | X |
| 16 | rs11864545 | 16 | | | | | | | | | | | |
| 17 | rs8133859 | Not found | | | | | | | | | | | |
| 18 | rs7907056 | 10 | COL13A1,N/A | | | | | | X | X | | | X |
| 19 | rs6055274 | 20 | | | | | | | | | | | |
| 20 | rs732985 | 1 | AL513218.1,ZFYVE9,CC2D1B | | | | | | X | | X | X | X |
| 21 | rs17052218 | 8 | AC120193.1 | | | | | | X | | X | | |
| 22 | rs12775535 | 10 | | | | | | | | | | | |
| 23 | rs10104915 | 8 | | | | | | | | | | | |
| 24 | rs35469947 | 1 | RAB4A,SPHAR | | | | | | X | | X | X | |
| 25 | rs7828453 | 8 | AC104248.1 | | | | | | X | | X | | |
| 26 | rs2919463 | 18 | | | | | | | | | | | |
| 27 | rs9498070 | 6 | | | | | | | | | | | |
| 28 | rs9828643 | 3 | CADPS | | | | | | X | | | | |
| 29 | rs16880351 | 5 | ITGA1 | | | | | | X | | | | |
| 30 | rs10174268 | 2 | CDKL4 | | | | | | X | X | | | |
| 31 | rs237157 | 16 | | | | | | | | | | | |
| 32 | rs8083143 | 18 | | | | | | | | | | | |
| 33 | rs3811010 | 1 | VANGL1 | | | | X | | | | | | X |
| 34 | rs16987299 | 19 | ZNF471 | | | | | | X | | X | | |

Figure S4: **NPDR gene regulation.** Top NPDR eQTLs associated with SCAI (suppressor of cancer cell invasion) RNA-Seq gene expression in major depressive disorder (MDD) study of 915 subjects. SNPs are tested genome-wide for NPDR-based association with SCAI. Tests are adjusted for MDD status and first 10 principal components. SNPs are ordered by NPDR P value, and annoation is provided for the SNPs chromosome location, nearest genes, and variant type.
